# A causal role for the right frontal eye fields in value comparison

**DOI:** 10.1101/2021.03.03.433716

**Authors:** Andres Mitsumasu, Ian Krajbich, Rafael Polania, Christian C. Ruff, Ernst Fehr

## Abstract

Recent studies have suggested close functional links between visual attention and decision making. This suggests that the corresponding mechanisms may interface in brain regions known to be crucial for guiding visual attention – such as the frontal eye field (FEF). Here, we combined brain stimulation, eye tracking and computational approaches to explore this possibility. We show that inhibitory transcranial magnetic stimulation (TMS) over the right FEF has a causal impact on decision-making, reducing the effect of gaze dwell time on choice while also increasing reaction times. We computationally characterize this putative mechanism by using the attentional drift diffusion model (aDDM), which reveals that FEF inhibition reduces the relative discounting of the non-fixated option in the comparison process. Our findings establish an important causal role of the right FEF in choice, elucidate the underlying mechanism, and provide support for one of the key causal hypotheses associated with the aDDM.

## Introduction

Despite the fact that decision-making and visual attention are both central features of cognition, we still know relatively little about how they interact. A prominent view in decision neuroscience is that the decision process consists of sequential sampling of information, with the choice implemented once the decision-maker accumulates enough net evidence in favor of one of the options (Ratcliff et al., 2016; Shadlen & Shohamy, 2016). Furthermore, experimental and theoretical accounts support the idea that such evidence accumulation is a domain-general mechanism underlying judgments about both the objective state of the physical world (perceptual decisions) (Bogacz et al., 2009; Forstmann et al., 2016; Gold & Heekeren, 2014; Hanks & Summerfield, 2017; O’Connell et al., 2018) and the subjective reward value of different choice options (Basten et al., 2010; Bhatia, 2013; Clithero, 2018; De Martino et al., 2013; Diederich, 2003; Fudenberg et al., 2018; Gluth et al., 2012; Hare et al., 2011; Hayden et al., 2011; Hunt et al., 2012; Hutcherson et al., 2015; Krajbich et al., 2015; Mormann et al., 2010; Philiastides & Ratcliff, 2013; Polania et al., 2014; Rodriguez et al., 2014; Roe et al., 2001; Tajima et al., 2016; Trueblood et al., 2014; Webb, 2018; Woodford, 2014; Zhao et al., 2020).

Visual attention allows us to selectively process the information in our environment, allocating greater computational resources to elements of interest in the visual scene, at the cost of diminishing the processing of unattended components (Carrasco, 2011; Chelazzi et al., 2011; Failing & Theeuwes, 2018; Itti & Koch, 2001). Thus, the sequential orienting of attention towards different stimuli is crucial for understanding the whole visual scene and guiding our gaze through it (Eimer, 2014). Attention can either be directed overtly (i.e., by eye fixation) or covertly (during constant fixation), but the effects on neural processing and the underlying causal mechanisms are thought to be strongly related (Moore & Zirnsak, 2017).

Investigations into the link between attention and decision making have shown that, during decision-making, subjects shift their gaze between the options until one of them is selected. These findings have led to proposals that overt visual attention may influence the evidence comparison process that guides choice behavior (Amasino et al., 2019; Ashby et al., 2016; Cavanagh et al., 2014; Folke et al., 2016; Hunt et al., 2018; Ian Krajbich et al., 2010; Shimojo et al., 2003; Towal et al., 2013). This proposed mechanism has been formalized by the attentional drift diffusion model (aDDM; (Fisher, 2017; Konovalov & Krajbich, 2016; Krajbich & Rangel, 2011; Krajbich et al., 2010; Krajbich, 2019; Smith & Krajbich, 2019; Tavares et al., 2017) and is supported by several reports of reliable correlations between gaze patterns and choice (Kovach et al., 2014; Smith & Krajbich, 2018; Stewart et al., 2015; Vaidya & Fellows, 2015) as well as experimental manipulations of attention that affect choice (Armel et al., 2008; Colas & Lu, 2017; Gwinn et al., 2019; Lim et al., 2011; Milosavljevic et al., 2012; Pärnamets et al., 2015), but see (Ghaffari & Fiedler, 2018; Newell & Le Pelley, 2018). However, most of the existing studies have manipulated attention by means of salient stimulus changes or direct experimental instruction, which may possibly induce experimenter demand effects on choice.

In the present study, we investigated the possible neural interface between attention and value-based choice using non-invasive brain stimulation, which offers a unique opportunity to test the functional contributions of brain areas reported to guide visual attention, without altering the experimental setup. We employed transcranial magnetic stimulation (TMS) and targeted the right human Frontal Eye Field (FEF) – a brain region often reported to be activated during the control of both eye movements and selective visual attention (Corbetta & Shulman, 2002; Hung et al., 2011; Juan & Muggleton, 2012; Marshall et al., 2015; Moore & Zirnsak, 2017; Schall, 2015) (Methods). We combined TMS with a value-based choice task, eye tracking, and the aDDM, to investigate how neural excitability modulations in the FEF affect both overt visual attention and the variables underlying the value-based choice process. Our choice task capitalizes on an important feature of the aDDM, namely that overt gaze has an amplifying effect on subjective values and so has a stronger effect on decisions between high-value items than low-value items (Shevlin & Krajbich, 2020; Smith & Krajbich, 2019; Westbrook et al., 2020). Thus, we employed a task with two overall-value conditions (low-value and high-value). This allowed us to test whether the FEF plays a causal role in the value-comparison process, and whether this effect is indeed stronger for higher-valued items, thereby indicating value amplification rather than just an attentional bias.

## Materials and methods

### Participants

Forty-five right-handed subjects (20 females, mean age ± SD = 23.14 ± 2.39) without a history of implanted metal objects, seizures or any other neurological or psychiatric disease participated in the experiment. No power analysis was used but the sample size was based on comparable TMS and eye-tracking studies at the time of data collection. Only subjects who reported not being on a diet were allowed to participate. Subjects were informed about all aspects of the experiment and gave written informed consent. Subjects received monetary compensation for their participation, in addition to receiving - at the end of the experiment - the chosen food item from a randomly selected choice trial. The experiments conformed to the Declaration of Helsinki and the Ethics Committee of the Canton of Zurich approved the experimental protocol.

### Experimental design

In a first task, subjects rated 148 food items (average duration of 10 minutes and 16 seconds, SD = 1 minute and 5 seconds). Every food item was presented individually on a computer screen for 2 seconds, followed by a rating screen (free response time). Subjects were instructed to press the space bar for those food items that they did not like at all, and to rate the remaining items on a scale from 0 to 10 based on how much they would like to eat that food at the end of the experiment. This rating task gave us a measure of the subjective value for each food item and allowed us to exclude disliked items (Fig. 1a).

**Figure 1.**
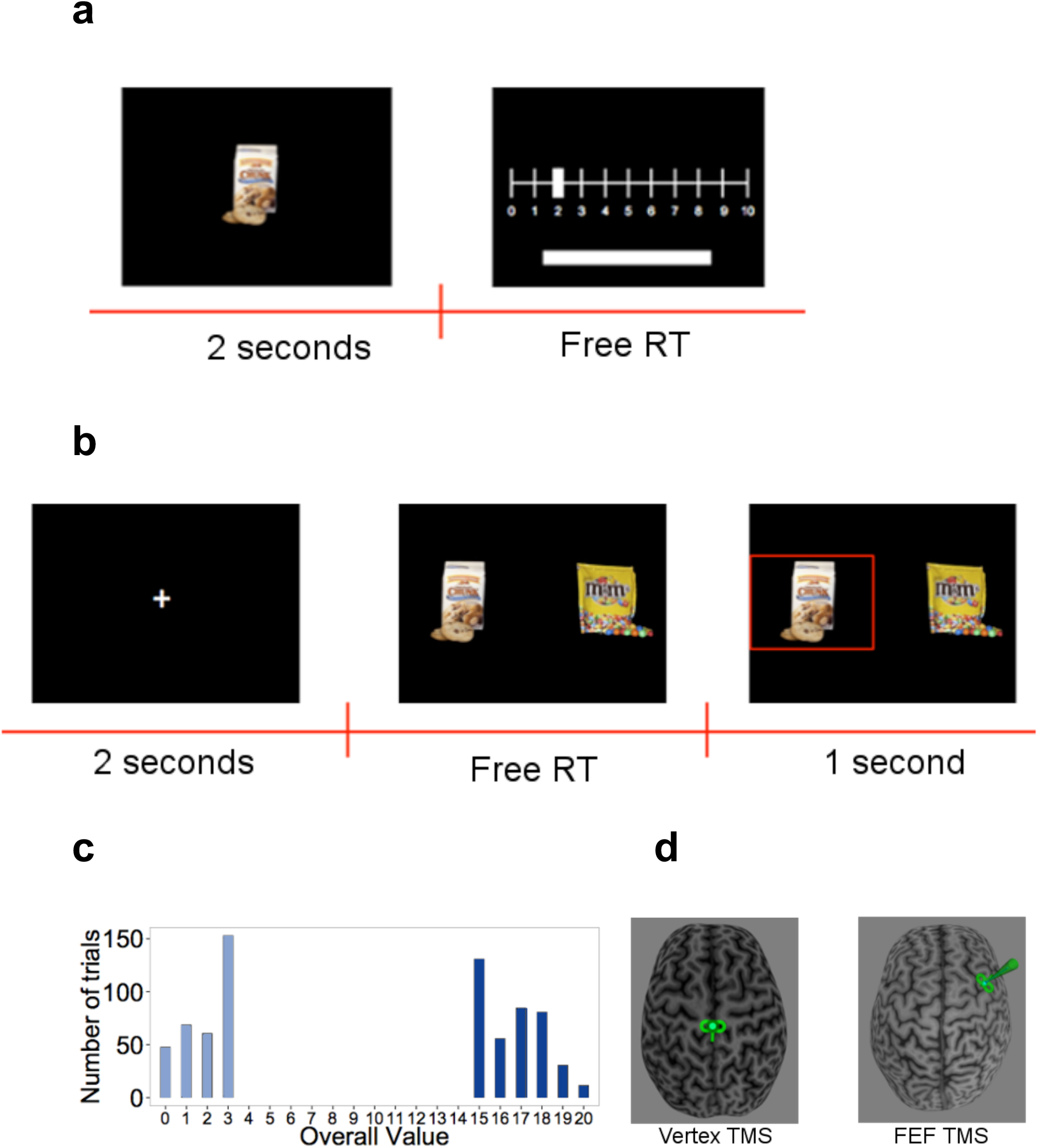
Experiment setup. **(a)** Rating-task timeline: Subjects saw each food item for 2 s and then rated how much they would like to eat it on a scale from 0 to 10, or excluded the item by pressing the space bar (no time limit). **(b)** Choice-task timeline: Subjects first had to fixate a central cross for 2 s. They then had to choose between the two presented food items using the keyboard. The chosen food was then highlighted for 1 s. **(c)** Histogram of overall value (OV) in the choice task: Trials were constructed to have either very high or low OV. **(d)** Stimulation: After the rating task, subjects received continuous theta-burst TMS over the vertex (left panel) or right FEF (right panel), depicted here schematically by the small green TMS coil symbol over one subject’s brain reconstruction.

After the rating task, subjects received inhibitory TMS (see below) on the right Frontal Eye Field or control stimulation on the vertex. We chose the right Frontal Eye Fields because this structure is one part of the well-established “dorsal attention network” (Corbetta & Shulman, 2002) and is known to contain neurons involved in target discrimination, saccadic eye movements, and covert attention towards specific visual field locations (Moore & Zirnsak, 2017; Schall, 2015). Moreover, FEF neurons have been shown to encode the reward value of objects in the current visual scene (Ding & Hikosaka, 2006; Glaser et al., 2016; Roesch & Olson, 2007; Serences, 2008) and are modulated in their attention-guiding function by dopaminergic neuromodulation (Noudoost & Moore, 2011). This suggests close functional interactions of the FEF with value-coding and dopaminergic reward systems, making this region an ideal candidate to house brain mechanisms responsible for top-down influences of attention on value computations during goal-directed choice. Due to the well-established right-hemispheric dominance for the orienting of spatial attention in humans (Heilman & Van Den Abell, 1980; Mesulam, 1981; Ruff et al., 2009), we chose the right FEF as our target region.

Subjects were randomly assigned either to the control stimulation group or the experimental stimulation group (FEF-TMS) before showing up to the experiment. Subjects were blind to their stimulation site, but the experimenters were not (at any part of the experiment or analysis). In the second task, immediately after the stimulation procedure, participants made 180 decisions between pairs of positively rated food items (average duration of 20 minutes and 36 seconds, SD = 5 minutes and 18 seconds) (Fig. 1b). The food items we presented were selected such that – for each participant – the difference in ratings between the left and right items (VD = left item value - right item value) was constrained to be -1, 0 or +1. This was done to focus on difficult choice problems where attention is most likely to affect the choice outcomes.

The task had two conditions, differing with respect to overall value (OV = left item value + right item value). In the high OV condition, subjects had to choose between two very appetitive (highly rated) foods, whereas decisions in the low OV condition only involved slightly appetitive (low rated) options (Fig. 1c).

During the binary choice task, we recorded subjects’ gaze at 250 Hz with an EyeLink-1000 (http://www.sr-research.com/). Choice trials with no gaze time on any food item were excluded from the analysis (0.008% of the pooled data from the 45 subjects). The mean (s.e.m.) number of trials dropped per subject was 1.44 ± 0.67. Both tasks were programmed in Matlab 2013b (Matworks), using the Psychophysics Toolbox extension (Brainard, 1997). We used R for statistical analysis as well as the HDDM analysis package for diffusion modeling (Wiecki et al., 2013).

### Transcranial magnetic stimulation

Subjects performed the binary choice task after receiving continuous theta burst TMS (cTBS) on the FEF (experimental group, n = 23) or the Vertex (control group, n = 22). In this TMS protocol – known to reduce neural excitability in the targeted area for up to 30 minutes - 600 magnetic pulses are administered over 40 seconds in bursts of 3 pulses at 50 Hz (20ms) repeated at intervals of 5 Hz (200 ms) (Huang et al., 2005). TMS pulses were delivered using a biphasic repetitive stimulator (Superapid2, Magstim, Withland, UK) with a 70 mm diameter eight-figure coil, and stimulation intensity was calibrated, for each subject, at 80% of active motor threshold. Prior to the experimental tasks, a structural T1-weighted anatomical MRI scan was acquired for every subject and reconstructed in 3-D for online neuro-navigation and precise placement of the TMS coil, with the Brainsight system (Rogue research, Montreal, Canada).

To stimulate the right FEF, the center of the coil was located on the right hemisphere, just anterior to the junction between the pre-central sulcus and superior frontal sulcus (MNI coordinates: xyz = 35.6 (+/-2.5),3.9 (+/- 2.7), 64 (+/-2.3)). During the stimulation procedure, the coil was positioned tangential to the targeted site and kept steady with a mechanical arm, with its handle oriented ∼ 45° in a rostral-to-caudal and lateral-to-medial orientation (i.e., parallel to the central sulcus). For stimulation on the vertex, the procedure was as above except that the center of the coil was located over the central fissure, at the intersection of the left and right central sulci, with the handle pointing backwards (Fig. 1d).

### Computational modeling

Our theoretical framework is the attentional drift diffusion model (aDDM) (Krajbich et al., 2010), though we also demonstrate that our results are robust to the modeling framework. In the standard DDM, decision makers accumulate noisy evidence for the options until the net evidence reaches a predefined threshold. In a value-based DDM, the rate of evidence accumulation (“drift rate”) thus reflects the subjective values of the options. The aDDM extends this model by allowing the drift rate to change within a trial, based on which option is fixated. In particular, the model assumes that gaze amplifies the value of the fixated (relative to non-fixated) option, shifting the drift rate towards that choice option and increasing the likelihood that it is chosen. Because of this amplifying (multiplicative) mechanism in this model, gaze has a stronger effect on the drift rate for high-value options, leading to shorter RTs and a larger effect of dwell time on choice probability (Shevlin & Krajbich, 2020; Smith & Krajbich, 2019; Westbrook et al., 2020) (Fig. 2a-b).

**Figure 2.**
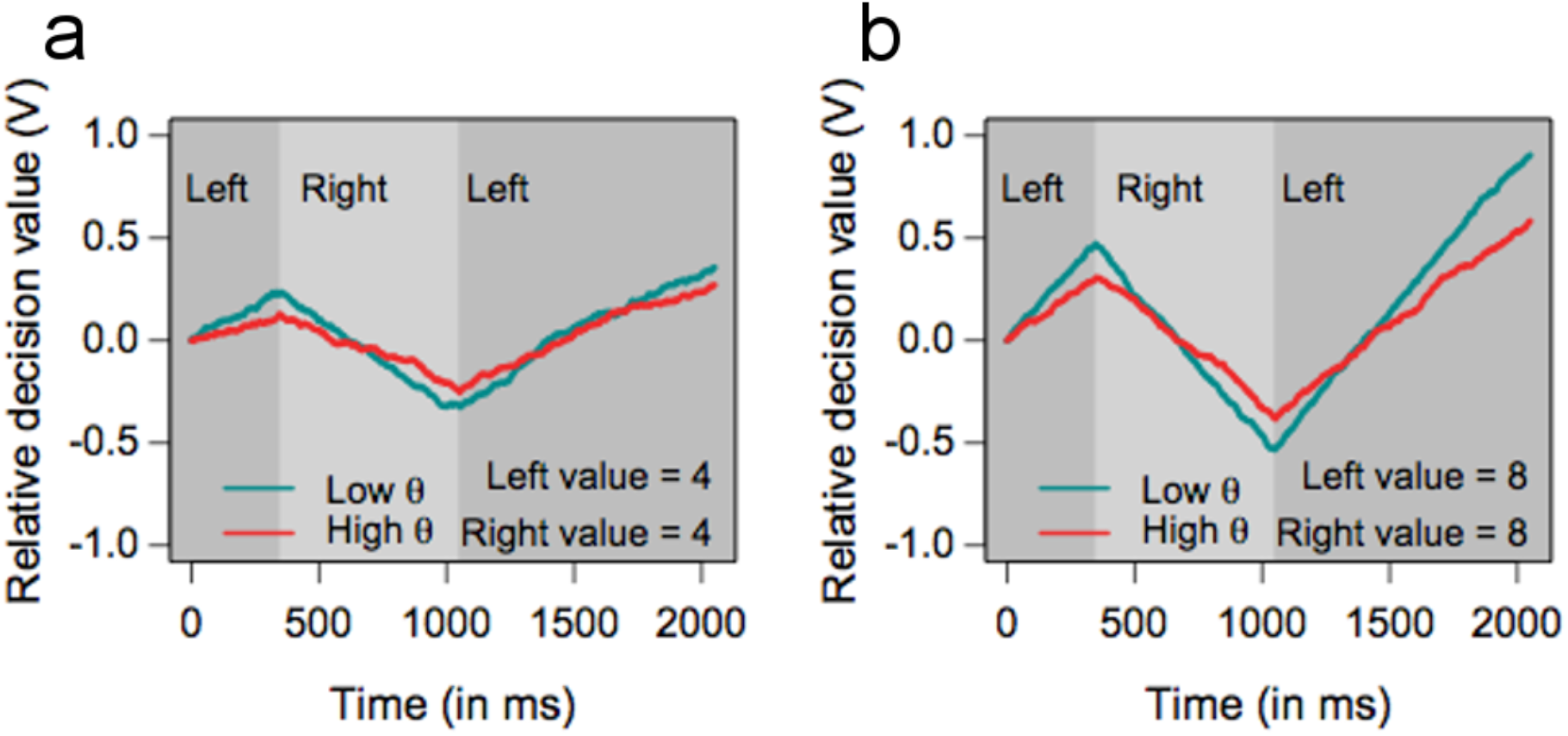
Simulations of the aDDM. To provide an intuition for why the aDDM makes different predictions for low and high OV, we simulated the aDDM, once with low values **(a)** and once with high values **(b)**. The simulations were run for a subject with a typical gaze discount factor (low θ-value, cyan line) (θ = 0.3) and a subject with less of a discount (red line) (θ = 0.5). Dark (light) gray areas indicate periods where the subject is looking at the left (right) item. The relative decision value (V) evolves over time with a slope that is biased toward the item that is being fixated. The left item is selected when V reaches 1 and the right item is selected when V reaches -1. The first thing to note is that holding OV constant (i.e. within a panel), a lower θ results in bigger changes in the slope (drift rate) when gaze shifts between Left and Right. The second thing to note is that holding θ constant, higher OV also results in bigger changes in drift rate when gaze shifts between Left and Right. Taken together, this means that the behavioral difference between low and high θ is much more pronounced for high vs. low OV. In particular, larger changes in drift rate lead to faster decisions and a stronger propensity to choose the longest attended option.

Formally, the aDDM captures the evidence accumulation process with a relative decision value (RDV) that evolves stochastically as follows. Let V_t_ be the value of the RDV at time t while d is a constant that controls the speed of change (in units of ms^−1^), and let r_left_ and r_right_ denote the values of the two options. Let θ (between 0 and 1) be a weight that discounts the value of the unattended alternative and, therefore, biases the RDV in favor of the attended one.ξ is white Gaussian noise with variance σ^2^, randomly sampled once every millisecond. Then, when a subject fixates on the left option, the RDV progresses according to

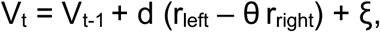

and when the subject fixates on the right option, the RDV changes according to

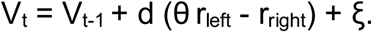

If the RDV reaches the +1 threshold the left reward is chosen and if it reaches the −1 threshold the right reward is chosen.

The parameter θ captures the degree to which the value of the fixated option is amplified by attention. A lower value of θ (closer to 0) indicates a stronger attentional influence, meaning that the decision maker is more likely to choose the longer-attended option and to respond faster (Fig. 2).

### Behavioral analysis and model fitting

We first investigated whether TMS had any effect on the gaze patterns in the data. We first conducted a logistic regression with clustered standard errors (mixed-effects models would not converge) in which first spatial fixation location (left =1, right = 0) was the dependent variable, regressed on OV interacted with a dummy variable for FEF-TMS group. We then conducted a mixed-effects logistic regression where first fixation location (coded as higher-rated item = 1, lower-rated item = 0) was regressed on OV interacted with a dummy variable for FEF-TMS group. Finally, we also conducted three regressions with clustered standard errors (mixed-effects models would not converge) to analyze the length of first, middle (all but first and last), and last dwell times. In each regression, log dwell time was regressed on TMS condition, position (left or right), OV, and value difference (VD) (left – right), all interacted together.

We next tested specific hypotheses about choices and RTs based on our theoretical framework. First, we conducted standard generalized linear model (GLM) analyses. Specifically, we conducted a trial-level logistic mixed-effects regression in which the chosen litem (left =1, right = 0) was the dependent variable, regressed on the value difference (left – right), a dummy variable for FEF-TMS group, a dummy variable for the OV condition, dwell-time advantage (total gaze dwell time spent on left – right item), and the interactions between the last three variables. The dwell-time advantage is our variable of interest as it captures the effect of increased attention on choice; value difference is included as a control variable to account for the difficulty of the decision. The model with full random effects would not converge, so we report the model with a random intercept and random slopes for value difference and dwell-time advantage; we obtain nearly identical results if we instead simply omit the random intercept from the full model. Additionally, we report the same logistic regression model but with clustered standard errors, which is an alternative way to account for repeated observations within subjects (Table S1).

To examine differences in RTs, we first computed median RTs in high and low OV trials for each subject and then compared them using paired t-tests. For a more sensitive analysis, we additionally performed non-parametric Kolmogorov-Smirnov tests between the pooled RT distributions.

Second, to test for differential effects in how TMS affected attention-based choice mechanisms, we utilized the hierarchical drift diffusion model (HDDM) package (Wiecki et al., 2013). This package uses Bayesian methods to estimate both group-level and subject-level DDM parameters. The package has a very useful regression feature, allowing the user to regress DDM parameters on trial-level features. A recent paper noted that one can use this feature to estimate gaze effects on drift rate (Cavanagh et al., 2014). A simple trick allows us to actually recover the attentional discounting parameter θ directly using this technique. In particular we run the following regression:

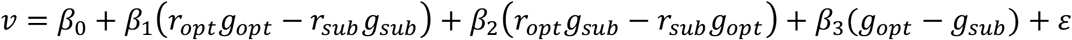

Where *v* is the drift rate, *r* are the values of the options, *g* is the fraction of the trial spent looking at the option, and the subscripts *opt* and *sub* indicate the higher and lower rated options, respectively. In the case of trials with equal-value options, we randomly assigned one of them to be *opt* and the other to be *sub. β*_1_ = *β*_2_ is the special case where gaze has no effect on drift rate; in that case the model reduces to simply *v* = *β*_*0*_ + *β*_1_ (*r*_*opt* −_ *r*_*sub*_) + ε which is the standard DDM. When *β*_1_ > *β*_2_, gaze has an amplifying effect on drift rate, as predicted by the aDDM, and *β*_2_*β*_1_ = θ. We include the third term (*β*_3_) to account for any possible additive effects of gaze on choice (Cavanagh et al., 2014; Westbrook et al., 2020).

## Results

### Does TMS affect the orienting of overt visual attention?

Before turning to the behavioral results, we first investigate whether FEF TMS had any effect on the gaze patterns, i.e., the deployment of overt attention. Any such effects would need to be accounted for in our subsequent analyses. Our subjects had a strong tendency to look first at the left food item (69.3%, 95% CI = [62.4%, 76.1%], p = 10^−6^) (Fig. 3). A logistic regression revealed that this tendency was reduced for high compared to low OV trials (OV: *β* = –0.194, CI = [–0.297, –0.091], p = 0.0002) in the Vertex group, but that this difference between OV conditions was eliminated in the FEF group (FEF x OV: *β* = 0.222, CI = [0.058, 0.385], p = 0.008) (Table S2).

**Figure 3.**
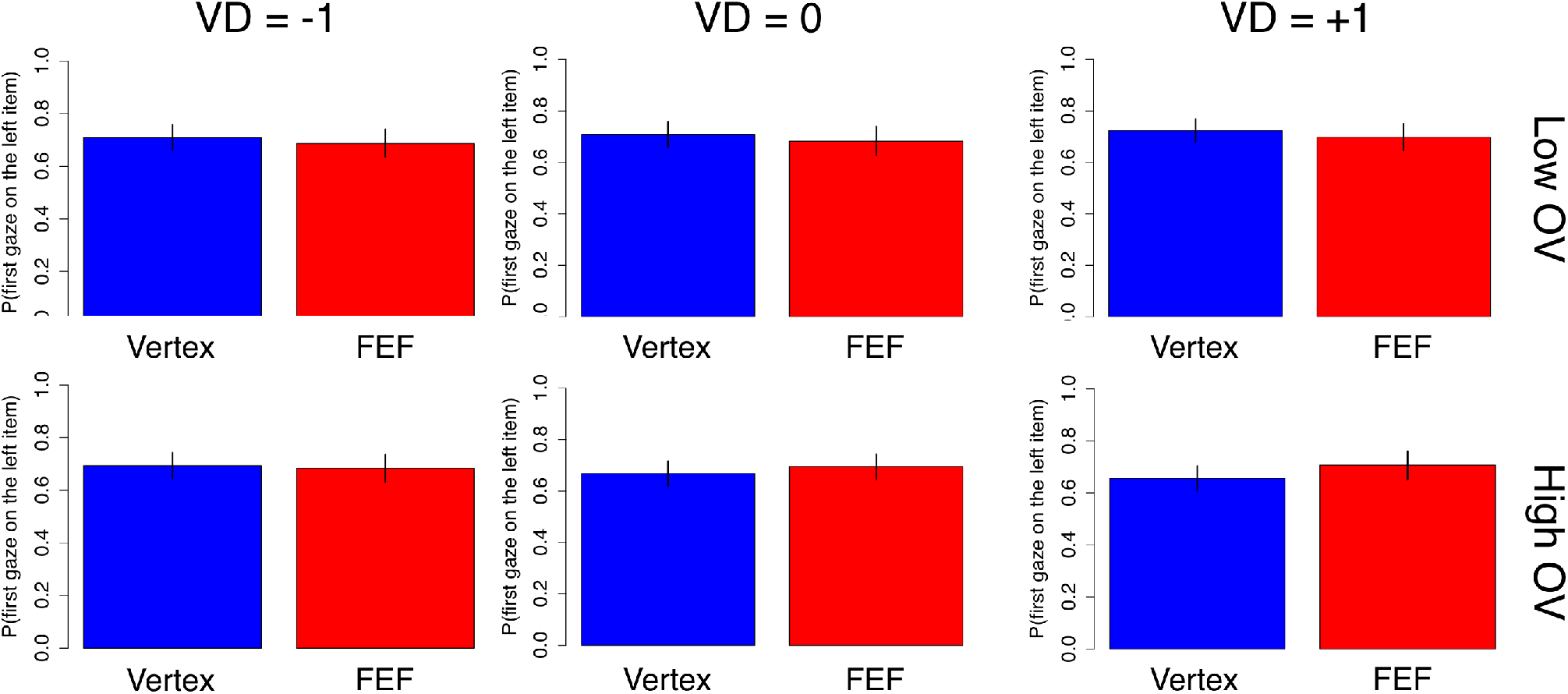
FEF effects on gaze patterns. Probability of looking first at the left food item during a trial. Subjects had a strong tendency to look left first. This tendency was slightly reduced in high vs. low OV trials for the Vertex group (in blue) but not the FEF group (in red). Bars are s.e.m.

A potential explanation for this effect is that for high OV trials, subjects are less likely to look left first because they are more likely to first look at the better item. We tested this idea with another logistic (mixed-effects) regression, looking at the probability of fixating the higher-rated item first. Consistent with prior work (Krajbich et al., 2010), a subject’s first gaze was no more likely than chance to go to the higher-rated item (50.2%, CI = [49.0%, 51.3%], p = 0.78). Moreover, we found no evidence for any OV or TMS effects. If anything, the probability of fixating the better item was numerically lower for high OV trials in the Vertex group (OV: *β* = –0.100, CI = [–0.253, 0.053], p = 0.2) and this effect was abolished under FEF TMS (FEF x OV: *β* = 0.126, CI = [–0.089, 0.340], p = 0.25) (Table S3).

In sum, the only effect of FEF TMS on first fixation location seems to be that it keeps subjects looking left first at the same rate in high (69.5%) and low (69.0%) OV trials, compared to Vertex TMS, which slightly reduces the rate of looking left first in high (68.4%) vs. low (71.4%) OV trials. We account for these effects in our later modeling to ensure that what we observe in the model parameters is not due to these differences in initial gaze allocation.

We also examined whether FEF TMS affected the gaze dwell times. In keeping with prior work (Krajbich et al., 2010, 2012), we separately analyzed first, middle, and last dwells. The basic reasoning is that first dwells cannot yet incorporate information about the non-fixated option, while subsequent dwells can. These first dwells also tend to be shorter than the rest. The last dwell of the trial is also different from the rest in that it is cut short by the crossing of the decision threshold.

In the regression analyses, we found no hint of any simple TMS effects or interaction effects (all p > 0.2) (Tables S4-6). Therefore, we conclude that FEF TMS did not induce any observable changes in dwell times relative to Vertex TMS.

### How does TMS affect choices and reaction times?

To test the aDDM predictions about how the right FEF should contribute to value-based choices, we compared trials with high or low OV, since the aDDM predicts stronger attentional effects (and therefore TMS modulation) for trials with high OV. Based on our theoretical framework, we tested several specific hypotheses with regard to choices and reaction times (RTs).

First, the aDDM predicts that subjects should select the longest attended alternative, and this effect should be stronger for high-OV trials. If the right FEF play a role in bringing about this effect, then the FEF (compared to Vertex) group should show a weaker effect of gaze on choice, particularly for the high-OV trials. We tested this hypothesis with a trial-level logistic mixed-effects regression in which the choice of the left item was the dependent variable, as a function of VD, TMS condition, OV, dwell-time advantage, and the interactions between the last three variables. The results confirmed all the predictions. During low-OV trials, subjects in the Vertex group were more inclined to choose the left food item as its dwell-time advantage increased (dwell-time advantage: *β* = 1.033, CI = [0.551, 1.515], p = 10^−5^). This effect increased strongly for high-OV trials (OV*dwell-time advantage: *β* = 0.625, CI = [0.376, 0.873], p = 10^−6^; Fig. 4a). The FEF group showed no difference to the Vertex group for the dwell-time-effect in low-OV trials (FEF x dwell-time advantage: *β* = 0.174, CI = [–0.489, 0.837], p = 0.61) but showed a substantial decrease relative to the Vertex group in high-OV trials (FEF x OV x dwell-time advantage: *β* = –0.709, CI = [–1.019, –0.400], p = 10^−5^), In fact, the difference in dwell-time effects between low and high OV was eliminated in the FEF subjects (Fig. 4a, Table S1).

**Figure 4.**
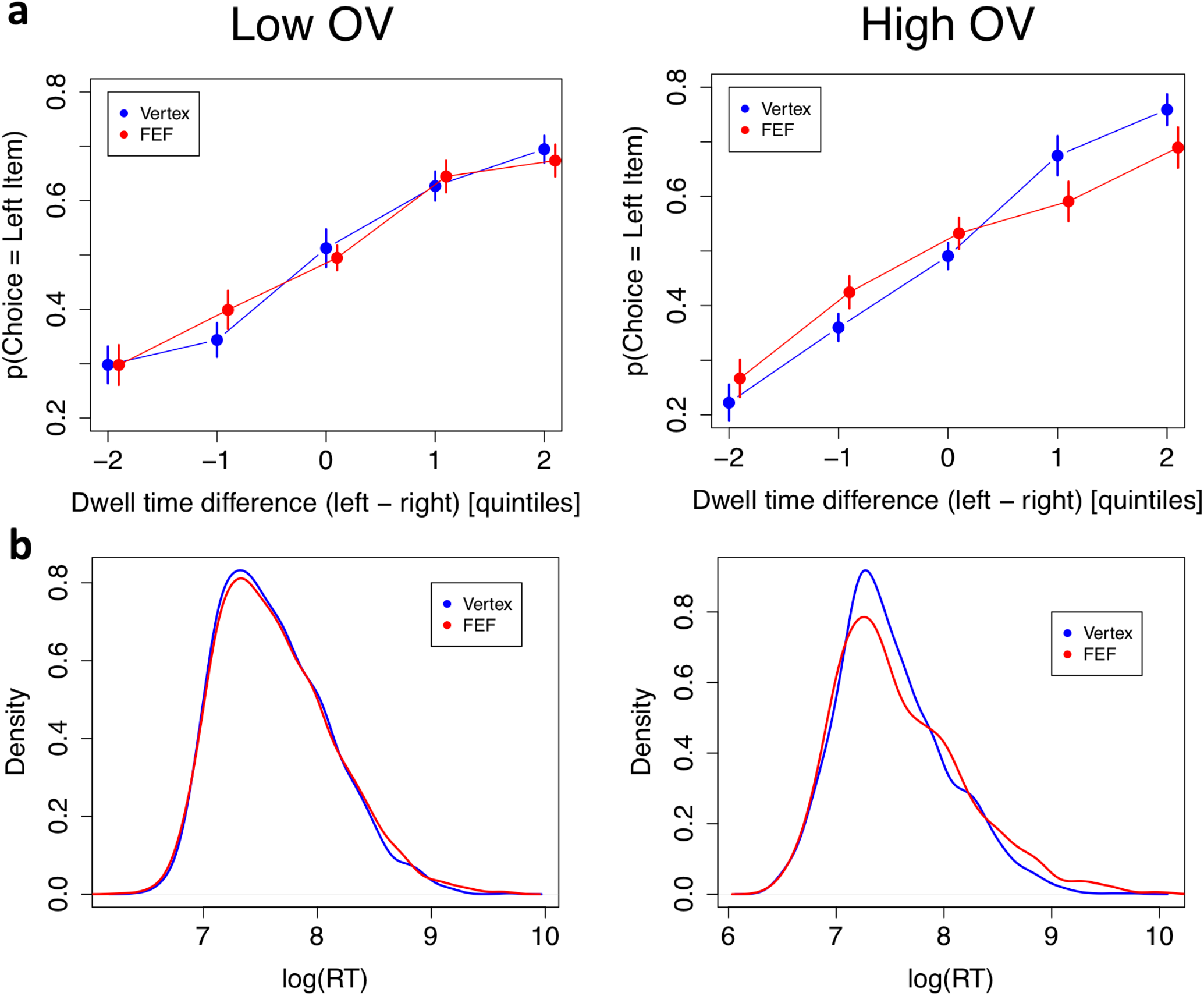
Behavioral results. The left panels are for low OV trials, the right panels are for high OV trials, and Vertex subjects are displayed in blue, FEF subjects in red. **(a)** Choice data: The probability of choosing the left item as a function of the total dwell time difference between the left and right items. Quintiles were determined at the subject level. Quintile 0 represents decisions where both items had similar total dwell times. Negative quintiles indicate more dwell time for the right item and positive quintiles indicate more dwell time for the left item. Bars are s.e.m. **(b)** Density plots of log RT.

Second, the aDDM also predicts that subjects should have shorter RTs for high-OV trials compared to low-OV trials. Again, this effect should be reduced for the FEF (relative to Vertex) group if FEF TMS reduces the attentional value-discounting effect on the choice process. We tested this with a trial-level mixed-effects regression in which log(RT) was the dependent variable, as a function of |VD|, TMS condition, OV, and the interaction between the last two variables. We included |VD| as a measure of decision difficulty, as is standard; here the effect was only marginal (|VD|: *β* = -0.02, CI = [–0.046, 0.007], p = 0.15) presumably because of the very narrow range of |VD| in our task. As expected, subjects in the Vertex group were faster in high-OV compared to low-OV trials (OV: *β* = -0.1, CI = [–0.149, –0.052], p = 0.0006), and this was not changed by TMS for low-OV trials (TMS: *β* = 0.021, CI = [– 0.136, 0.178], p = 0.79). The expected interaction between TMS and high-OV was not significant, but pointed in the correct direction and numerically cut the effect of OV on RT by about half (TMS x OV: *β* = 0.06, CI = [–0.019, 0.139], p = 0.15).

For a more sensitive analysis of the RTs, we utilized the Kolmogorov-Smirnov (K-S) method to test for differences between the RT distributions. K-S tests revealed no difference between stimulation groups for low-OV trials (D = 0.022, p = 0.71) but a significant difference for high-OV trials (D = 0.064, p = 0.005) (Fig. 4b). They also revealed significant differences between high and low-OV trials in both groups, though the difference was numerically larger for the Vertex group (D = 0.097, p = 10^−8^) than for the FEF group (D = 0.091, p = 10^−7^). Admittedly, these tests do not account for repeated measures per subject, so they should be treated with caution.

### Model fitting

According to our hypothesis, inhibitory TMS on the right FEF should decrease the gaze discount (i.e., increase θ) on the unattended option during the evidence accumulation process. The behavioral analyses reported above are consistent with this hypothesis. Next, we made simultaneous use of choice and RT data, using diffusion modeling, to provide more direct evidence for this hypothesis, by showing that the estimated θ parameters were indeed higher for subjects in the FEF-TMS group compared to the Vertex-TMS group.

To do so, we used HDDM to fit a hierarchical diffusion model that accounts for the effects of gaze on choice. In addition to the standard threshold separation (*a*) and non-decision time (*T*_*er*_) parameters, we also estimated drift rate as a function of the food ratings and dwell-time proportion (see Methods).

Looking first at threshold separation, we found no difference between the Vertex group (*a* = 2.63, CI = []) and the FEF group (*a* = 2.68, CI = [2.43, 2.93]) (t(40.2) = 0.30, p = 0.77). This clearly indicates that there was no change in response caution between the two groups. Looking next at the non-decision-time parameter, we again found no difference between the Vertex group (*T*_*er*_ = 664 ms, CI = [611, 718]) and the FEF group (*T*_*er*_ = 695 ms, CI = [637, 752]) (t(42.9) = 0.76, p = 0.45). This indicates that there was also no change in general RT components that are separate from the decision process. In the model, we also accounted for potential additive effects of gaze on choice (i.e., changes that do not reflect modulation of value evidence but that are constant and independent of item values). We did observe significant additive effects of gaze direction, but importantly, these effects did not significantly differ between the Vertex group (*β*_3_ = 0.551, CI = [0.415, 0.688]) and the FEF group (*β*_3_ = 0.695, CI = [0.48, 0.91]) (t(37.1) = 1.10, p = 0.28).

Having established that FEF TMS does not affect general, value-independent response processes, we turn to our key test. Comparing the two groups on the θ estimates we found that the estimated θ was significantly lower for the Vertex group (θ = 0.776, CI = [0.708, 0.844]) than for the FEF group (θ = 0.937 CI = [0.818, 1.055]) (t(34.9) = 2.30, p = 0.03). This demonstrates that the effects on choice behavior induced by TMS on the right FEF were specifically due to the multiplicative aDDM effects, and not due to changes in response caution, non-decision-time, or additive gaze effects.

It is important to note that the results that we have presented demonstrate that FEF- TMS reduces the effect of gaze on choice even when the observed gaze patterns are accounted for. Thus, FEF-TMS has effects on value discounting during choice that are independent of any effects the stimulation may have on gaze patterns themselves.

## Discussion

Based on our computational modeling framework, we developed a paradigm that allowed us to causally manipulate value-based choice by inhibiting the right FEF. Formal computational modeling using HDDM confirmed that the sole effect of the FEF inhibition was to reduce attentional effects on choice, i.e., to increase the gaze discount factor θ. Importantly, this change was measured after accounting for the limited effects of the FEF stimulation on the gaze patterns themselves as well as on value-independent decision processes.

Our results provide a neural validation for a central assumption in the aDDM, namely that attention amplifies the subjective values of the items, leading to larger effects on choice for higher-valued items (Smith & Krajbich, 2019). In designing our experiment, we capitalized on this feature of the model by contrasting low- and high-value trials. By showing that FEF stimulation affected choices and RTs in high-value but not low-value trials, we confirmed a role for right FEF in attention-based value modulation during choice, and further validated the multiplicative nature of the aDDM (Westbrook et al., 2020).

More generally, our results build on a literature documenting the DDM-like neural mechanisms underlying value-based choice. These papers have used EEG (Polania et al., 2014), fMRI (Basten et al., 2010; Gluth et al., 2012; Hare et al., 2011; Lim et al., 2011; Rodriguez et al., 2015) and their combination (Pisauro et al., 2017) to identify neural signatures of evidence accumulation in structures such as the dorsal/posterior medial prefrontal cortex, dorsolateral prefrontal cortex, and intraparietal sulci. Our results suggest that the right FEF may play a critical role in modulating the resulting activity in this set of regions, through its effects on attention.

Our results also suggest some interesting parallels in how attention may operate, by means of FEF and its feedback projections, in both perceptual and value systems in the human brain. As for perception, previous psychophysical studies have shown that, relative to unattended visual stimuli, perceptual sensitivity is enhanced for attended elements of the visual scene (Barbot et al., 2011; Herrmann et al., 2010; Montagna et al., 2009; Pestilli et al., 2007; Pestilli & Carrasco, 2005). These investigations suggest that attended visual stimuli may have stronger neural representations than unattended ones, and this view has been supported by neurophysiological reports in humans (Kastner et al., 1998; Liu et al., 2005; O’Craven et al., 1997) and non-human primates (Connor et al., 1997; Martínez-Trujillo & Treue, 2002; Reynolds et al., 2000; Reynolds & Desimone, 2003). More precisely, these investigations have shown that neuronal populations that code for attended locations of the visual scene display enhanced activity, relative to neurons representing unattended locations. Importantly, the FEF is a possible source of these attention-dependent modulations: Investigations combining brain stimulation with neuroimaging techniques have shown that modulations of neuronal activity in the FEF induce top-down modulatory effects on both behavior and neuronal activity in early visual areas that resemble effects of attention (Moore & Armstrong, 2003; Moore & Fallah, 2004; Ruff et al., 2006; Silvanto et al., 2006). Additionally, it has been suggested that attention-dependent behavioral effects are due to a boost in synchronization between FEF and V4 at the gamma-band frequency (Gregoriou et al., 2009).

In light of this information, it is interesting that TMS of the FEF also leads to decreasing modulatory effects of attention on the value of non-attended items. Could these behavioral effects reflect that the FEF may exert similar top-down modulatory effects on value representations? Extensive research indicates that such value representations are found in a large network of brain areas including the ventromedial prefrontal cortex (vmPFC), orbitofrontal cortex (OFC), ventral striatum (vStr) and the posterior parietal cortex (PPC) (Bartra et al., 2013; Boorman et al., 2009; Clithero & Rangel, 2014; Cromwell & Schultz, 2003; Kahnt et al., 2014; Knutson et al., 2001; Padoa-Schioppa & Assad, 2006; Plassmann et al., 2007; Platt & Glimcher, 1999). Moreover, these value-related BOLD signals are increased when attention is directed to the value (rather than other aspects) of objects (Grueschow et al. 2015; Grabenhorst & Rolls 2008), or when participants attend to specific value-relevant aspects of a given stimulus (Lim et al., 2011). Interestingly, in another analogy to the perceptual domain, attending to the value of items leads to increases in fronto-posterior synchronization in the gamma frequency (Polania et al., 2014), and decreasing the degree of this coherence by brain stimulation leads to inaccurate value-based choices (Polanía et al., 2015). None of these studies have explicitly examined to what degree these attentional effects on value-based choices may involve the FEF, but our current results suggest that this should be a fruitful area for future studies.

Taking together, our findings demonstrate the relevance of the FEF for attention-dependent modulations of value-based decision processes, and they suggest directions for future investigations on the interaction between visual-attention brain networks and areas coding value-signals.

## Acknowledgements

Ernst Fehr acknowledges support from the European Research Council (Advanced Grant 295642-FEP). Christian Ruff acknowledges support from the Swiss National Science Foundation (SNF grant 105314_152891) and the European Research Council (Consolidator Grant COG-2016-725355). Ian Krajbich acknowledges support from the National Science Foundation (NSF Career Award 1554837) and the Cattel Sabbatical Fund. We thank the staff at the SNS lab and the Neuroscience Center Zurich ZNZ for practical support.

## Author contributions

A.M., C.C.R., and I.K. designed the experiment with input from R.P. and E.F. A.M. carried out the experiment. A.M. and I.K. carried out the behavioral analysis. I.K. performed model fits. A.M., I.K., C.C.R. and E.F. wrote the manuscript.

---

**Table S1.** Choice behavior. Logistic mixed-effects regression for the probability of choosing the left item. The baseline condition in this regression is Vertex stimulation with low OV decisions.

|  | Estimate | Standard Error | Z value | P value |
| --- | --- | --- | --- | --- |
| Intercept | -0.027 | 0.056 | -0.49 | 0.63 |
| **VD** | **0.637** | **0.049** | **13.12** | **10<sup>-16</sup>** |
| TMS (FEF = 1, Vertex = 0) | 0.029 | 0.078 | 0.37 | 0.71 |
| **Dwell-time advantage (ms)** | **1.033** | **0.246** | **4.20** | **10<sup>-5</sup>** |
| OV condition (high = 1, low = 0) | -0.002 | 0.070 | -0.02 | 0.98 |
| TMS x dwell-time advantage | 0.174 | 0.338 | 0.51 | 0.61 |
| TMS x OV condition | -0.019 | 0.098 | -0.19 | 0.85 |
| **OV condition x dwell-time advantage** | **0.625** | **0.127** | **4.93** | **10<sup>-6</sup>** |
| **TMS x OV condition x dwell-time advantage** | **-0.709** | **0.158** | **-4.49** | **10<sup>-5</sup>** |

**Table S2.**
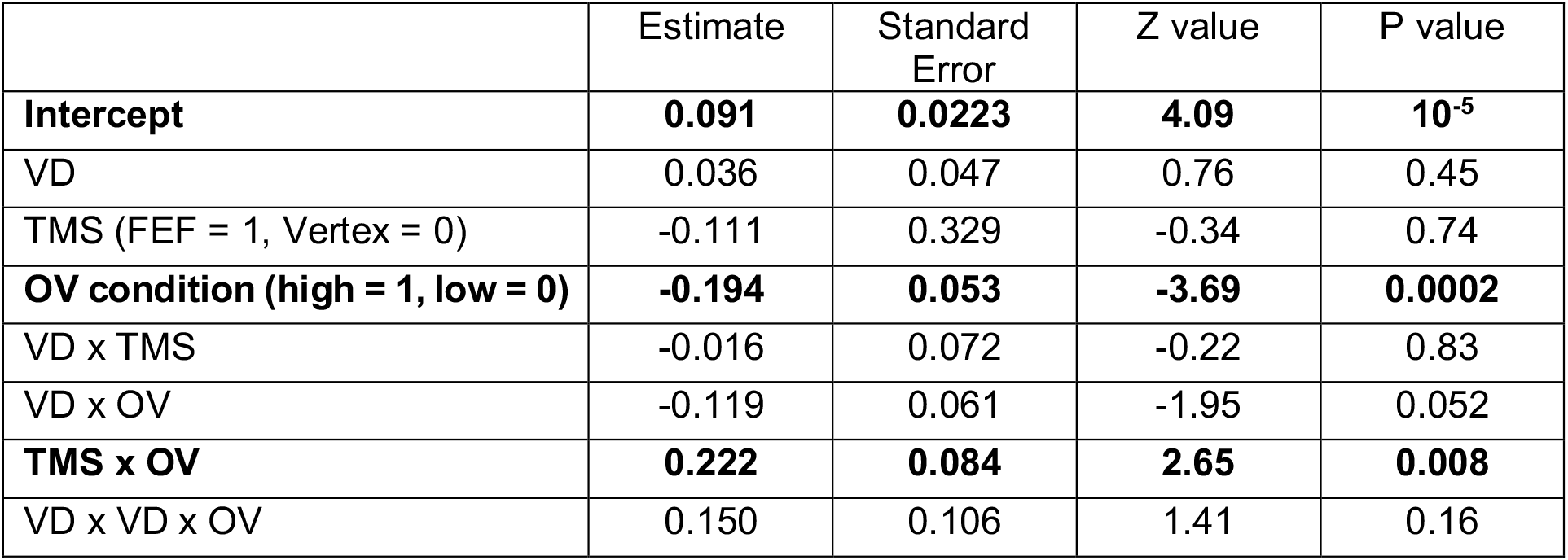
First gaze left. Logistic regression for the probability that the first gaze is to the left item. Clustered standard errors were used to account for repeated observations within-subject. The baseline condition in this regression is Vertex stimulation with low OV decisions.

**Table S3.** First gaze to better item. Logistic mixed-effects regression for the probability that the first gaze is to the better item. The baseline condition in this regression is Vertex stimulation with low OV decisions. For this analysis we excluded trials with VD equal to 0 because then there is no better food item. The model includes full random effects at the subject level.

|  | Estimate | Standard Error | Z value | P value |
| --- | --- | --- | --- | --- |
| Intercept | 0.029 | 0.055 | 0.52 | 0.60 |
| TMS (FEF = 1, Vertex = 0) | -0.011 | 0.077 | -0.15 | 0.88 |
| OV condition (high = 1, low = 0) | -0.100 | 0.078 | -1.29 | 0.20 |
| TMS x OV condition | 0.126 | 0.109 | 1.15 | 0.25 |

**Table S4.**
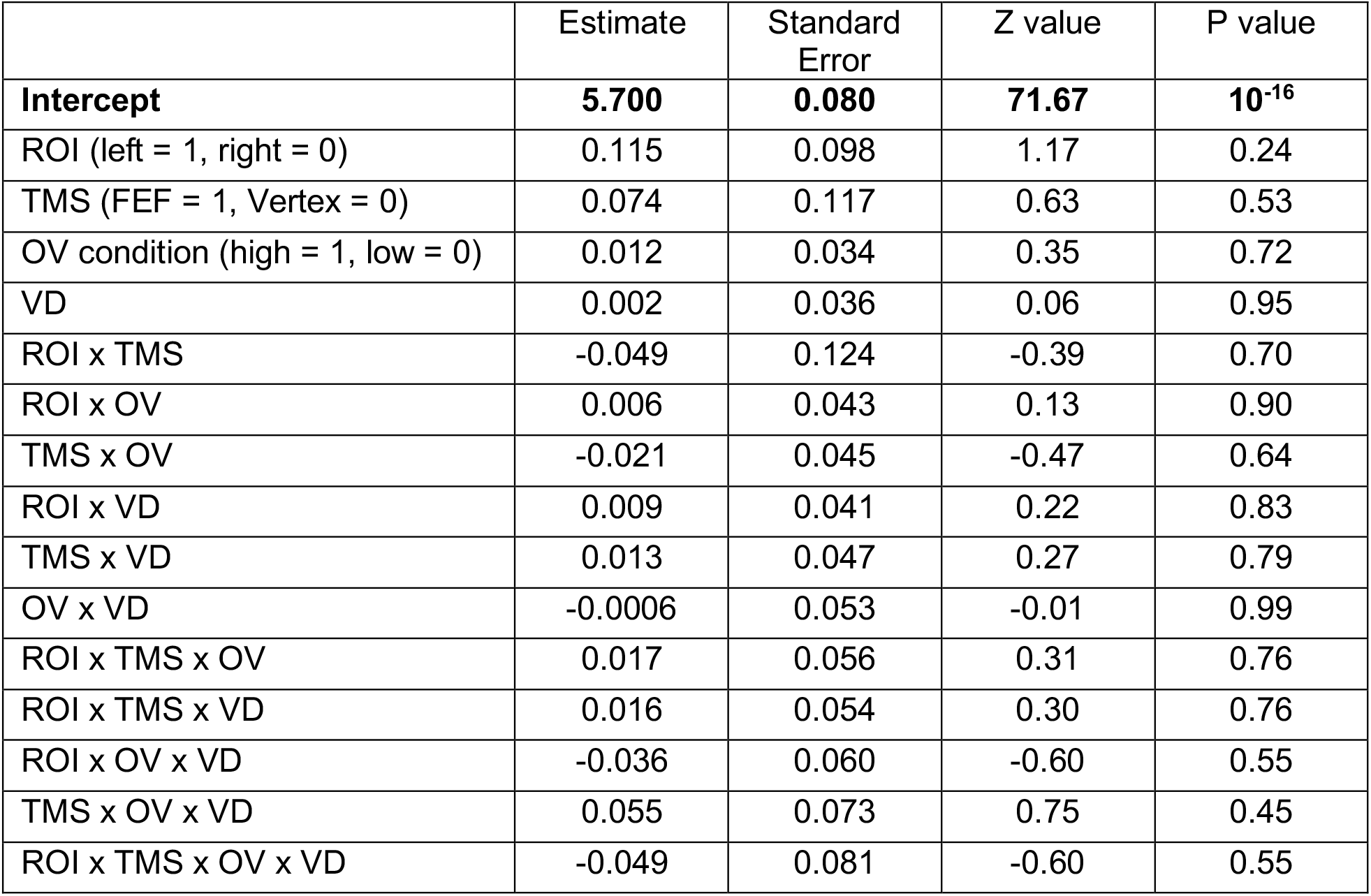
First gaze dwell time. Linear regression for the first gaze log(dwell time). Clustered standard errors were used to account for repeated observations within-subject. The baseline condition in this regression is Vertex stimulation with low OV decisions.

|  | Estimate | Standard Error | Z value | P value |
| --- | --- | --- | --- | --- |
| **Intercept** | **5.700** | **0.080** | **71.67** | **10<sup>-16</sup>** |
| ROI (left = 1, right = 0) | 0.115 | 0.098 | 1.17 | 0.24 |
| TMS (FEF = 1, Vertex = 0) | 0.074 | 0.117 | 0.63 | 0.53 |
| OV condition (high = 1, low = 0) | 0.012 | 0.034 | 0.35 | 0.72 |
| VD | 0.002 | 0.036 | 0.06 | 0.95 |
| ROI x TMS | -0.049 | 0.124 | -0.39 | 0.70 |
| ROI x OV | 0.006 | 0.043 | 0.13 | 0.90 |
| TMS x OV | -0.021 | 0.045 | -0.47 | 0.64 |
| ROI x VD | 0.009 | 0.041 | 0.22 | 0.83 |
| TMS x VD | 0.013 | 0.047 | 0.27 | 0.79 |
| OV x VD | -0.0006 | 0.053 | -0.01 | 0.99 |
| ROI x TMS x OV | 0.017 | 0.056 | 0.31 | 0.76 |
| ROI x TMS x VD | 0.016 | 0.054 | 0.30 | 0.76 |
| ROI x OV x VD | -0.036 | 0.060 | -0.60 | 0.55 |
| TMS x OV x VD | 0.055 | 0.073 | 0.75 | 0.45 |
| ROI x TMS x OV x VD | -0.049 | 0.081 | -0.60 | 0.55 |

**Table S5.** Middle gaze dwell time. Linear regression for the middle gaze log(dwell time). Clustered standard errors were used to account for repeated observations within-subject. The baseline condition in this regression is Vertex stimulation with low OV decisions.

|  | Estimate | Standard Error | Z value | P value |
| --- | --- | --- | --- | --- |
| **Intercept** | **6.382** | **0.054** | **118.84** | **10<sup>-16</sup>** |
| ROI (left = 1, right = 0) | -0.022 | 0.043 | -0.51 | 0.61 |
| TMS (FEF = 1, Vertex = 0) | 0.022 | 0.086 | 0.26 | 0.80 |
| **OV condition (high = 1, low = 0)** | **-0.064** | **0.021** | **-3.08** | **0.002** |
| **VD** | **-0.039** | **0.016** | **-2.43** | **0.02** |
| ROI x TMS | 0.056 | 0.054 | 1.04 | 0.30 |
| ROI x OV | -0.029 | 0.023 | -1.25 | 0.21 |
| TMS x OV | 0.041 | 0.032 | 1.28 | 0.20 |
| **ROI x VD** | **0.063** | **0.029** | **2.19** | **0.03** |
| TMS x VD | 0.008 | 0.024 | 0.33 | 0.74 |
| OV x VD | 0.039 | 0.023 | 1.70 | 0.09 |
| ROI x TMS x OV | 0.044 | 0.042 | 1.06 | 0.29 |
| ROI x TMS x VD | 0.023 | 0.040 | 0.58 | 0.56 |
| ROI x OV x VD | -0.064 | 0.035 | -1.84 | 0.07 |
| TMS x OV x VD | 0.029 | 0.037 | 0.78 | 0.43 |
| ROI x TMS x OV x VD | -0.038 | 0.057 | -0.65 | 0.51 |

**Table S6.**
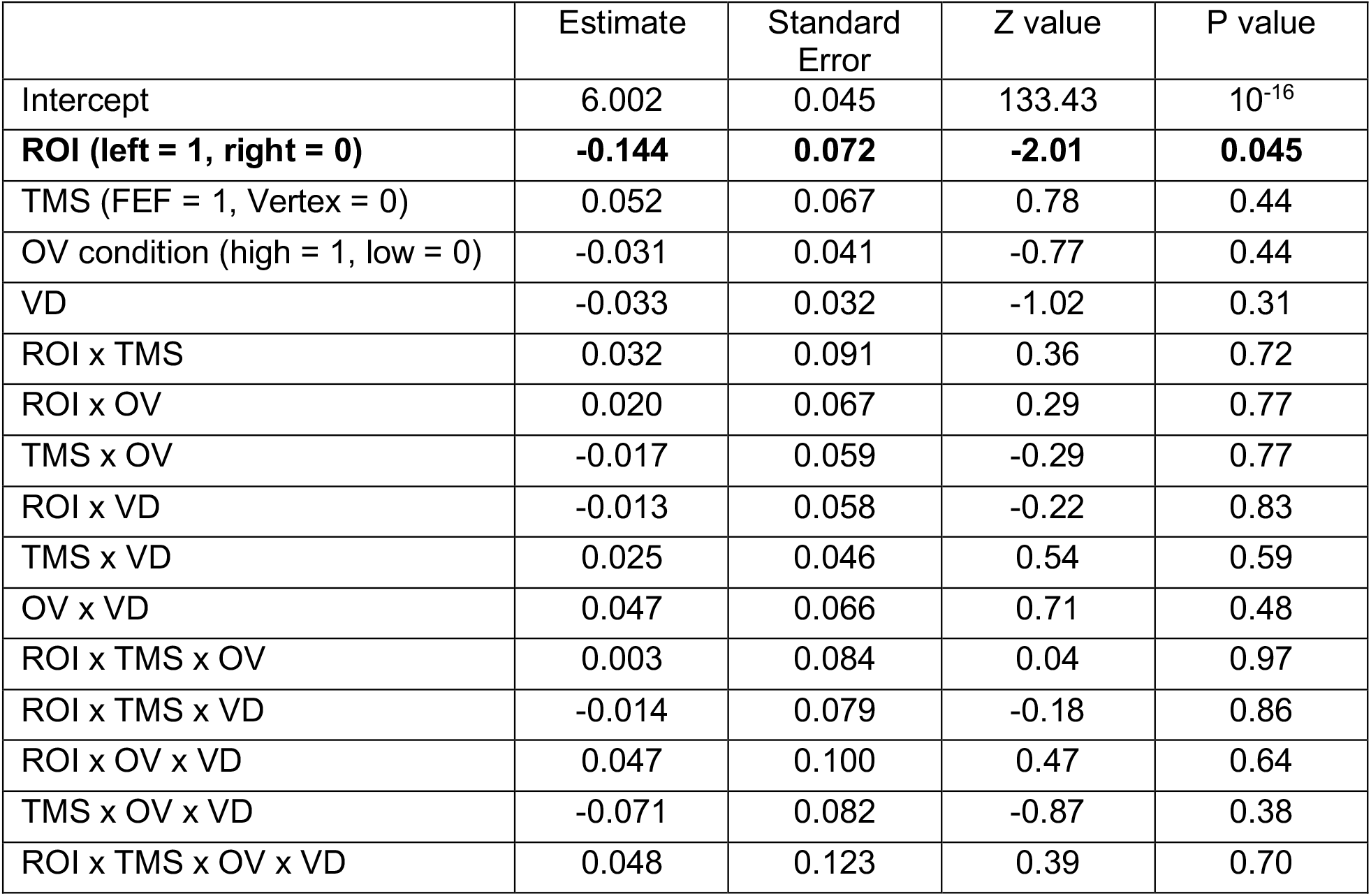
Last gaze dwell time. Linear regression for the last gaze log(dwell time). Clustered standard errors were used to account for repeated observations within-subject. The baseline condition in this regression is Vertex stimulation with low OV decisions.

